# Uncovering Xenobiotics in the Dark Metabolome using Ion Mobility Spectrometry, Mass Defect Analysis and Machine Learning

**DOI:** 10.1101/2021.11.21.469449

**Authors:** MaKayla Foster, Markace Rainey, Chandler Watson, James N. Dodds, Facundo M. Fernández, Erin S. Baker

## Abstract

The identification of xenobiotics in nontargeted metabolomic analyses is a vital step in understanding human exposure. Xenobiotic metabolism, excretion, and co-existence with other endogenous molecules however greatly complicate nontargeted studies. While mass spectrometry (MS)-based platforms are commonly used in metabolomic measurements, deconvoluting endogenous metabolites and xenobiotics is often challenged by the lack of xenobiotic parent and metabolite standards as well as the numerous isomers possible for each small molecule *m/z* feature. Here, we evaluate the use of ion mobility spectrometry coupled with MS (IMS-MS) and mass defect filtering in a xenobiotic structural annotation workflow to reduce large metabolomic feature lists and uncover potential xenobiotic classes and species detected in the metabolomic studies. To evaluate the workflow, xenobiotics having known high toxicities including per- and polyfluoroalkyl substances (PFAS), polycyclic aromatic hydrocarbons (PAHs), polychlorinated biphenyls (PCBs) and polybrominated diphenyl ethers (PBDEs) were examined. Initially, to address the lack of available IMS collision cross section (CCS) values for per- and polyfluoroalkyl substances (PFAS), 88 PFAS standards were evaluated with IMS-MS to both develop a targeted PFAS CCS library and for use in machine learning predictions. The CCS values for biomolecules and xenobiotics were then plotted versus *m/z*, clearly distinguishing the biomolecules and halogenated xenobiotics. The xenobiotic structural annotation workflow was then used to annotate potential PFAS features in NIST human serum. The workflow reduced the 2,423 detected LC-IMS-MS features to 80 possible PFAS with 17 confidently identified through targeted analyses and 48 additional features correlating with possible CompTox entries.

## Introduction

Measuring chemical exposure is extremely challenging due to the range and number of anthropogenic molecules encountered in our daily lives, as well as their complex biochemical transformations throughout the body. Metabolomic measurements host an abundance of exposure information as they are composed of xenobiotics originating from diet, lifestyle, and environmental exposure, in addition to information about endogenous molecules from primary pathways, secondary signaling, and microbial communities within our bodies (**Figure 1**).^1–4^ With all these considerations, millions of molecules are estimated in the human metabolome. These estimates include more than 16,000 known endogenous metabolites, 1500 drugs, 22,000 food constituents,^5^ and thousands of xenobiotics from environmental exposures since over 882,000 parent chemicals are present in the CompTox database alone,^6,7^ not even including all the xenobiotic degradants and metabolites. These xenobiotics and their degradants and metabolites are also included in the exposome, which was first introduced by Dr. Christopher Wild in 2005 and defined as “the totality of exposures from conception onwards”.^8,9^ Since xenobiotic exposure is known to affects a person’s health, and various epidemiological studies have revealed links between environmental exposures and many different infections, conditions, and diseases including various cancers and Alzheimer’s disease, evaluating these chemicals in metabolomic studies is essential.^10^

**Figure 1.**
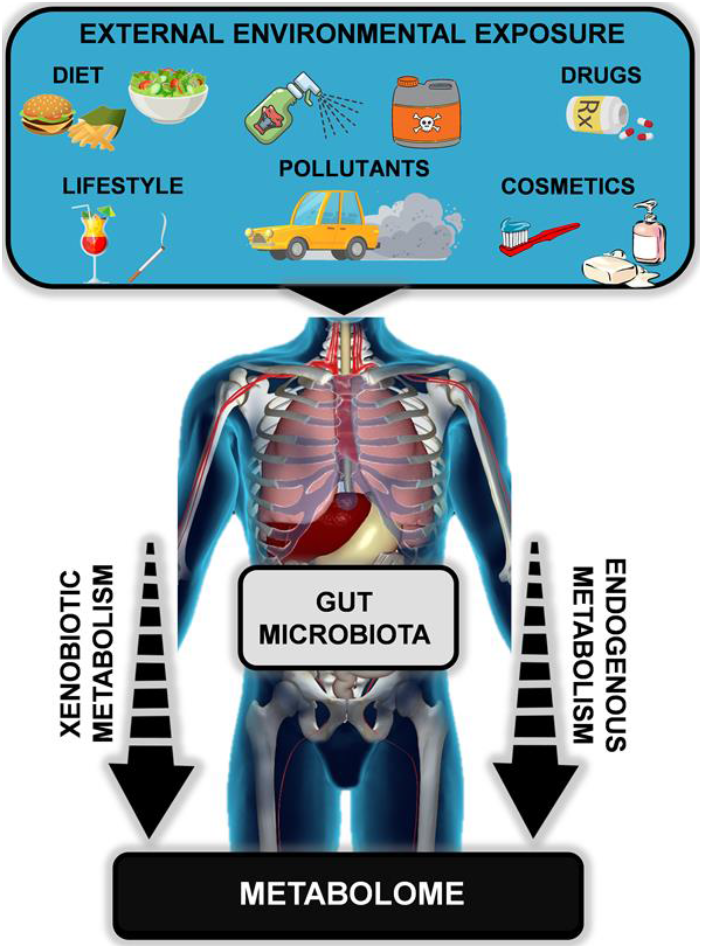
Molecular constituents in the metabolome. External environmental exposures, xenobiotic metabolism, endogenous metabolism, and gut microbiota all contribute to the metabolome.

To date, direct measurements of xenobiotics biofluids and tissues are performed to attain exposure data. Biological responses are also commonly used to infer the extent of chemical exposure as many xenobiotics can be cleared from the body prior to any observed perturbations. Omic-based measurements (e.g., genomics transcriptomics, proteomics, and metabolomics) are therefore extremely valuable for gaining insight into the molecular disruptions. While genomic, transcriptomic, and proteomic analyses have rapidly progressed over the last two decades,^1,2^ xenobiotic and endogenous metabolite measurements have not advanced to nearly as great a degree even though they are essential for the direct evaluations of chemical exposure.

To gain a better understanding of the molecules in the metabolome and exposome, numerous analytical strategies have been employed.^11^ Unfortunately despite advances in nuclear magnetic resonance (NMR) spectroscopy, chromatography, and mass spectrometry (MS), only ∼10% of the tens of thousands of small molecule features to date are reliably annotated.^2,12^ These unknown molecules, together with those that are not detected by the specific analytical platform used, are collectively known as the “dark metabolome”.^2^ Thus, our ability to better understand this unexplored fraction of the metabolome must improve in order to draw more informative connections between human health and the environment.^2,12^ Unfortunately there are many challenges involved in these measurements. For example, xenobiotic compounds in pesticides, cosmetics, and air pollutants can be taken into the body by a variety of avenues with the metabolome reflecting both chronic and acute exposure to such substances.^13,14^ Additionally, the various molecular properties of these compounds require both polar and nonpolar sample extractions, specific concentration steps, and analytical platforms having dual ionization methods and different chromatography types. Furthermore, the resulting complex datasets require alignment and mining by sophisticated computational tools, which often demand machine learning (ML) capabilities.^15–17^ However, as automated analytical capabilities and computational approaches advance, our ability to address these challenges continues to grow with each year.

Computational approaches are extensively leveraged to map xenobiotics within the complexity of the dark metabolome.^18,19^ Xenobiotics of great interest due to their known high toxicities include per- and polyfluoroalkyl substances (PFAS),^20,21^ polycyclic aromatic hydrocarbons (PAHs),^22,23^ polychlorinated biphenyls (PCBs)^24,25^ and polybrominated diphenyl ethers (PBDEs).^26–28^ Each of these classes includes a variety of subclasses and also unknowns from metabolism and degradation. The numerous resulting species therefore require progressively more sophisticated techniques to detect their presence, enable structural characterization, and evaluate their temporal evolution and distribution. Since many of these xenobiotic classes have characteristic molecular and structural traits, such as PFAS all sharing the presence of C-F bonds and PCBs having chlorinated biphenyl moieties, these can be exploited to increase detection specificity. For example, Kendrick mass defect (KMD) analysis has been important for distinguishing different molecular classes as it can probe repetitive patterns in complex datasets by normalization to specific atomic or functional group components. In the case of PFAS, KMD analysis with a CF_2_ repeating unit pinpoints molecules with and without CF_2_ functional groups.^20,21^ Furthermore, since PFAS have varying head groups and linkages including hydrocarbon regions (fluorotelomer sulfonate (FTSs)), ether bonds (PFECAs) and branched C-F backbones,^10,12^ subclass distinction is possible even within this single xenobiotic class.^29–35^ Orthogonal separation dimensions have also shown utility for PFAS analyses. In a study coupling liquid chromatography, ion mobility spectrometry and MS (LC-IMS-MS), the multidimensional evaluations uncovered specific PFAS trends as the *m/z* versus IMS collision cross section (CCS) plots distinguished each PFAS subclass studied.^36,37^ These CCS and *m/z* evaluations are progressively being utilized in more unknown small molecule identification efforts with specific utility in database matching.^36,38^ However if standards are not available, CCS prediction with ML as a structural filter is necessary and has been readily applied with several groups showcasing theoretical and experimental CCS value differences of less than 2%.^36,39,40^ As such, CCS prediction through ML has great promise in elucidating information for the many metabolites and xenobiotics without standards.^40,41^

In this manuscript, we present a xenobiotic structural annotation workflow that utilizes mass defects and CCS values to filter down large metabolomic feature lists and elucidate potential detected xenobiotic classes and species. Due to its recent linkages to significant health impacts, PFAS were of particular interest for this evaluation, however since only a few PFAS experimental CCS values were available, initially we evaluated 88 different PFAS standards with drift tube IMS. These values were utilized to create a targeted library that we uploaded to the free repository Panorama^42^ within the open-source software tool Skyline,^43^ so that it is available to the research community. Next, the PFAS CCS values and PCBs, PBDEs, and PAHs values available in the Unified CCS Compendium^44–46^ were inputted into the open-source ML CCS prediction tool, CCSP 2.0, to calculate theoretical CCS values since many standards are not available for xenobiotics in these classes. Finally, a xenobiotic annotation workflow utilizing the CCS values and mass defect filtering was applied to a nontargeted metabolomic feature list for NIST serum to discover potential PFAS.

## Materials and Methods

### Xenobiotic Chemical Standards and PFAS Serum Extraction

Eighty-seven PFAS standards were characterized with LC-IMS-MS to obtain CCS and *m/z* values (**Table S1**). Examples of the different PFAS subclasses studied include perfluoroalkyl carboxylic acids, perfluoroalkyl sulfonic acids, and Nafion byproducts. PFAS standards were obtained from Wellington, EPA, Chemours, SynQuest, 3M, Acros Organics, ASM, and Alfa Aesar (**Table S1**). Each standard was diluted in water (Thermo Fisher, Waltham, MA) or methanol (Thermo Fisher, Waltham, MA) to 5-10 µg mL^-1^ for LC-IMS-MS analyses and CCS IMS measurements, as outlined below. The dilution solvent was chosen based on the solubility of the target compound and the degradation properties of each PFAS standard (**Table S1**). For PAHs, PCBs and PBDEs, CCS values were obtained from the CCS Compendium^47^ and other literature sources.^46^

PFAS were extracted from NIST human serum 909c (Gaithersburg, MD). In this extraction, 50 µL of thawed serum was mixed with 1 µL of the heavy labeled PFAS mixture MPFAC-C-ES (Wellington Laboratories, ON, Canada) and 2 µL of the heavy labeled standard M3HFPO-DA or GenX (Wellington Laboratories, ON, Canada) for quantitation. The serum mixture was then vortexed for 30 s, followed by protein precipitation with 300 µL of cold (−20°C) acetonitrile (Thermo Fisher, Waltham, MA). The sample was vortexed again for 30 s and chilled for 30 min at −20°C, followed by centrifugation at 12,500 g for 5 min. After centrifugation, 200 µL of the acetonitrile supernatant was transferred to a microcentrifuge tube and placed in a SpeedVac until dry. The residue was then reconstituted in 100 µL of a 40% methanol (Thermo Fisher, Waltham, MA) and 60% water (Thermo Fisher, Waltham, MA) solution containing 2 mM ammonium acetate (Thermo Fisher, Waltham, MA). Finally, the sample was vortexed for 30 s and transferred to a LC vial for analysis. Blanks containing methanol were also used in the study to determine extraction and instrumental PFAS contaminates. All blanks underwent the same extraction procedure as the NIST serum.

### LC-IMS-MS Analyses

The reverse phase LC method utilized for measuring monoisotopic masses and CCS values for the PFAS standards and serum extracts has been previously described.^27^ Briefly, this method uses an Agilent 1290 Infinity LC system (Agilent Technologies, Santa Clara, CA) equipped with a C18 column (Agilent ZORBAX Eclipse Plus, 2.1 × 50 mm, 1.8 μm). The mobile phases utilized contained 18 MΩ cm water from an ELGA Purelab Flex purification system (High Wycombe, UK), ammonium acetate (Fisher Scientific, Waltham, MA), and/or LC-IMS grade methanol (Fisher Scientific, Waltham, MA). The composition of Mobile Phase A (MPA) was 5 mM ammonium acetate in water, while Mobile Phase B (MPB) consisted of 5 mM ammonium acetate 95% methanol and 5% water. The LC gradient was as follows (% MPB, Time (min)): 10%:0 min, 10%:0.5 min, 30%:2 min, 95%:14 min, 100%:16.5 min, followed by 6 min re-equilibration at 10% MPB.

For the IMS-MS analyses, an Agilent 6560 IM-QTOF MS (Agilent Technologies, Santa Clara, CA) was utilized in negative ion mode with an electrospray ionization (ESI) source (Agilent Jet Stream). Since negative mode analyses were performed, the deprotonated ion species of all PFAS were monitored, in addition to the [M-COOH]^-^ decarboxylated species for PFAS containing carboxylic acid functionalities such as the polyfluoroalkyl ether carboxylic acids and perfluoroalkyl carboxylic acids. Following ionization, the ions were pulsed into the drift tube, containing ∼3.950 Torr N_2_ gas every 60 ms, with 100 µs ion gating to minimize peak broadening and obtain the highest possible resolving power. An electric field of 17.2 V cm^-1^ was utilized in the drift tube and the TOF was operated in the 50-1700 *m/z* range. IMS-MS nested spectra were acquired using the MassHunter Acquisition Software B.09, and data were analyzed using IMS-MS Browser Version 10.0 preceding spreadsheet exportation (Microsoft Excel) for further data analysis. All ^DT^CCS_N2_ values were calculated using a single-field method that has been extensively tested by multiple laboratories.^27,33–36^ This method utilizes Agilent tune mix ions as IMS calibrants and triplicate CCS values are measured for the different ions in separate experiments over three days to confirm both method reproducibility and instrument stability. The CCS values collected with this method were all highly precise, with an RSD lower than 0.3%.

### Mass Defect and Kendrick Mass Defect Calculations

To evaluate the mass defect and KMDs for CF_2_ and CH_2_-containing species, **Equations 1** through **3** were utilized.^20–22,24^

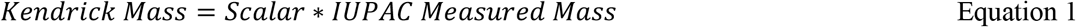

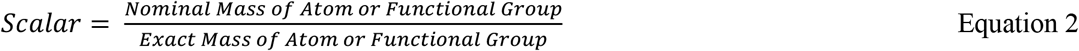

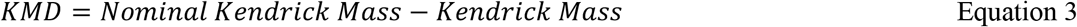

Since the scalar factor in Equation 2 is obtained as the nominal mass of the target compound divided by its exact mass, it varies for each repeating unit or functional group of interest. For example, for CF_2_ it is 50/49.9968 and for CH_2_ it is 14/14.0156 (showing the exact mass to 4 decimal points). Since mass defect is based on ^12^C, its scalar factor is equal to one since the nominal and exact masses of carbon are both defined as 12 Da (or 12/12). Additionally, all mass defects and KMDs are reported in parts per thousand (ppt) for optimal visualization.

### Machine Learning CCS Prediction

For the ML analyses, PFAS, PCBs/PBDEs, and PAHs were utilized to develop a Python-based ML CCS prediction approach for xenobiotics. A previous version of the CCS Predictor (CCSP)^48,49^ was used as a starting point, and new CCSP 2.0 code was prepared in JupyterLab V2.2.6 using Python V3.8.5 and the following dependencies: RDKit V2020.09.1, PubChemPy V1.0.4, Pandas V1.1.3, Numpy V1.19.2, Matplotlib V3.3.2, and Plotly V4.14.3. For the PFAS study set, 102 PFAS experimental CCS values obtained from drift tube IMS were utilized. Seventy-eight of these CCS values corresponded to deprotonated species and the remaining were for polyfluoroalkyl ether carboxylic acid and perfluoroalkyl carboxylic acid subclasses which can also be detected in their decarboxylated form ([M-COOH]^-^). For ML, all species were neutralized through proton addition in ChemDraw Professional V16.0 (Perkin-Elmer, Waltham, MA). The PBDEs/PCBs set consisted of 16 parent PBDEs, 20 hydroxide PBDE derivatives, 23 PCBs and 13 PCB hydroxide derivatives. The 72 CCS values were obtained from the Unified CCS Compendium,^44^ and included compounds detected in their [M-Cl+O]^-^ and [M-Br+O]^-^ forms. Again, all ions were neutralized using Chem Draw Professional. Finally, the PAH dataset consisted of 26 parent PAHs and 4 PAH ketone derivatives. Here 29 PAHs were obtained from the Unified CCS Compendium and one was retrieved from CCSBase.^50^ Each detected protonated PAH ion was neutralized through deprotonation prior to ML. Additionally, structural representations for each molecule were encoded as a MOL file based on its IUPAC International Chemical Identifier (InChI) using RDKit. When available, the InChI was obtained from the structure’s PubChem entry. Any structure without a PubChem entry was drawn in ChemDraw Professional and its InChI exported. For each InChI, 1613 2D molecular descriptors were calculated with the Python package Mordred V1.2.0.^51^ Evaluation of each ML algorithm randomly assigned half of the molecules in each set to a calibration (training) group for model construction and the other half were held for validation. Molecular descriptors remaining constant across all calibration sets were discarded and any descriptor not applicable to one or more molecules in the calibration or validation sets was excluded. The remaining descriptors were *z*-transformed using the mean and standard deviation of the descriptors in the calibration set.

The PFAS, PDBE/PCB and PAH sets were next used to train the support vector regression models employing a linear kernel. Two hyperparameters were tuned in model development: (1) the regularization parameter *C*, which determines the penalty gradient associated with non-zero residuals and (2) *ϵ*, the radius of an epsilon-tube in which no penalty is accrued for non-zero residuals. Hyperparameters were selected using a grid search approach, where *C* and *ϵ* were evaluated pairwise using 5-fold cross-validation. *C* was selected from nine options (0.015625, 0.03125, 0.0625, 0.125, 0.25, 0.5, 1, 2, 4), and *ϵ* from five options (0.01, 0.05, 0.1, 0.5, 1). The hyperparameter pair yielding the lowest root mean squared error of cross-validation (RMSECV) was chosen for further evaluation. Linear-SVR model construction was performed using the SVR module of the Python project SciKit-Learn V0.23.2 (scikit-learn.org), and the grid search used the GridSearchCV module of the same project. Feature selection was carried out using Recursive Feature Elimination (RFE) to reduce the feature (molecular descriptor) space. Features with the lowest model weight were iteratively removed until RMSECV increased and RFE was performed using the RFECV module of SciKit-Learn. The full CCSP 2.0 Jupyter notebook is publicly available at https://github.com/facundof2016/CCSP2.0.

## Results and Discussion

## CCS versus m/z Separation of Biomolecules and Xenobiotics

In this study, 88 PFAS standards were characterized with IMS-MS to evaluate their CCS values, mass defects and various KMDs (**Table S1**). These CCS values were utilized both to populate a targeted Skyline database and for ML prediction of theoretical CCS values since standards are not available for many of the possible 9000+ PFAS proposed by the EPA. The Skyline database can be found in **Table S2** and also online at https://panoramaweb.org/NCSU%20-%20Baker%20Lab/PFAS%20Library/project-begin.view.

Due to both deprotonation and decarboxylation of some PFAS species such as those for the polyfluoroalkyl ether carboxylic acid and perfluoroalkyl carboxylic acid subclasses, 102 unique PFAS CCS values were obtained from the IMS-MS analyses. CCS versus *m/z* plots for the measured PFAS showed linearity and distinction for the different subclasses as previously observed in a smaller PFAS study^36^ (**Figure 2A**). The trends were then further evaluated in relation to other biological and xenobiotic molecules by using CCS values from the Unified CCS Compendium^44^ for fatty acids, phosphatidylcholines, bile acids, nucleosides, nucleotides, carbohydrates, PCBs, PBDEs, and PAHs. Interestingly, the biomolecules had higher slopes in the CCS versus *m/z* plots than molecules with more halogen atoms such as the PFAS and PBDEs (**Figure 2B**). One exception was for the bile acids as they had a flatter slope than all the other biomolecules studied since they have a common steroid backbone, but because they were initially shifted up in the graph due to the steroid atomic composition, they a higher CCS per *m/z* relationship compared to the halogenated molecules. The PCB trend was also notable as PCBs with fewer chlorines were initially observed between the biological and halogenated regions, however as the number of chlorines grew, they dipped into the halogenated region. Upon investigation, the observed separation between biomolecules and halogenated xenobiotics was attributed to the fact that halogens have a much larger mass than the hydrogens in biomolecules, but their atomic size is not as massive. Thus, when a hydrogen is replaced by a halogen, a large *m/z* change is noted, but the molecular size change is not of the same magnitude. For example, a 1 Da hydrogen atom replaced by a 19 Da fluorine atom greatly increases the *m/z* of the molecule but does not significantly increase its structural size as the van der Waals radius of fluorine is ∼1.47 Å while hydrogen is ∼1.20 Å.^52^ This effect emphasizes the power of utilizing IMS-MS for the detection of xenobiotics as it allowed clear separation of halogenated species, which is particularly powerful in PFAS and lipids studies as they both can be extracted from the same sample and ionized in negative ESI mode.

**Figure 2.**
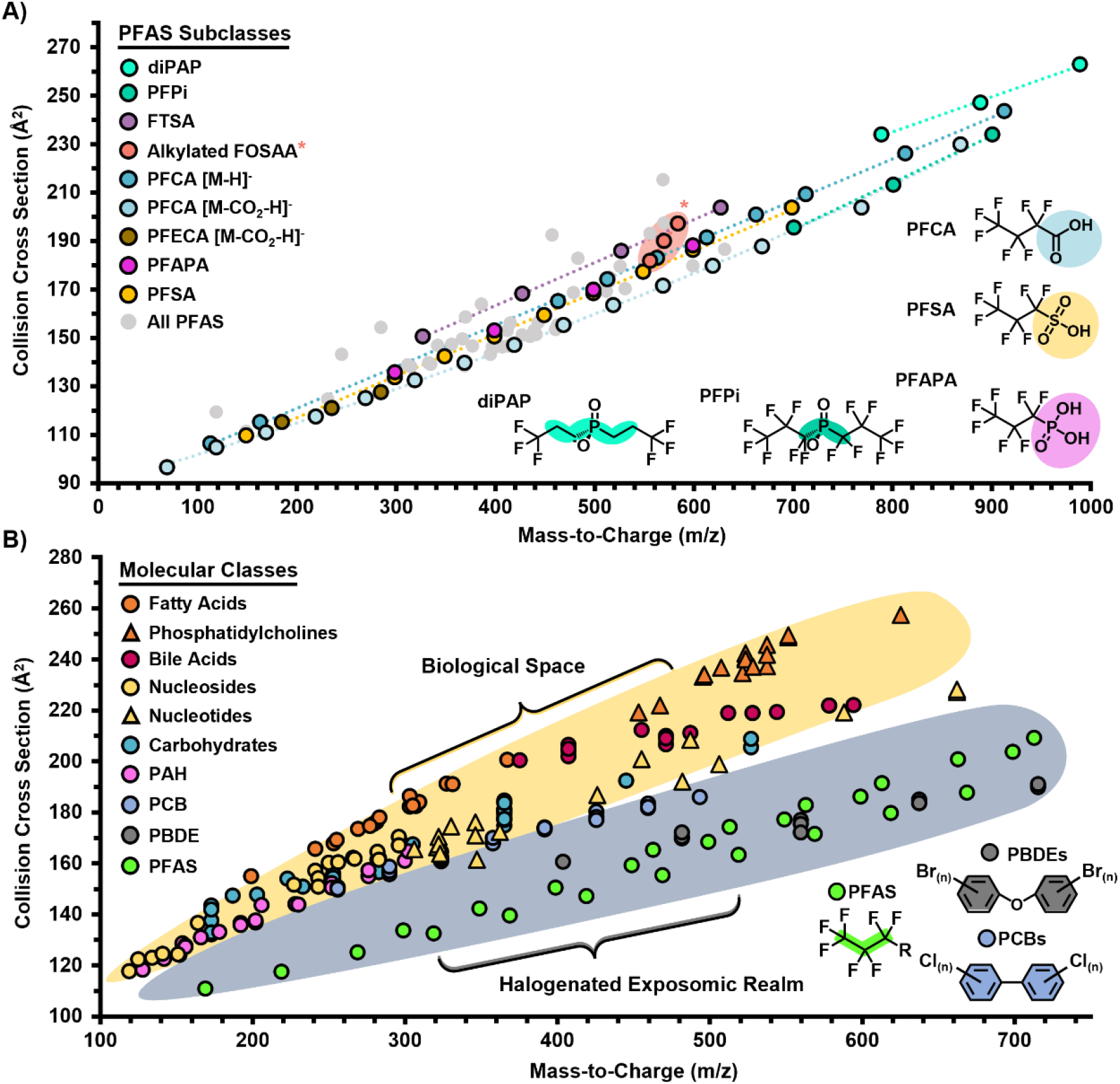
Mass-to-charge versus ion mobility size separations for classes and subclasses of xenobiotics and biomolecules. **A)** In an investigation of the 88 PFAS studied, trend lines were noted for each of the distinct PFAS subclasses (*e*.*g*., FTSA, PFCA and PFSA) allowing their identification and separation by IMS-MS. Several PFAS subclasses are highlighted to illustrate their main structural characteristics and trends. **B)** Biomolecules and heavily halogenated xenobiotics also showed clear distinction in their *m/z* versus CCS plots due to the greatly increased m/z of halogens versus hydrogen, but small size difference. All noted CCS values are ^DT^CCS_N2_.

### Utilizing CCS, Mass Defect, KMD and m/z for PFAS Identification

Mass defect and KMD analyses were next performed for 201 PFAS frequently found in the environment and of particular interest to the Environmental Protection Agency (EPA) (**Figure 3**). Since evaluation of mass defects only requires accurate *m/z* values (see Equations in the Methods section), no IMS experimental measurements or standards were necessary. The 290 resulting values shown in **Table S3** are based on the evaluation of both deprotonated and decarboxylated ions for PFAS containing a carboxylic acid headgroup. Since PFAS have such a high number of fluorine atoms and CF_2_ repeat units, KMD-CF_2_ values were studied first. In the analyses of *m/z* versus KMD-CF_2_ space, a slope of zero was observed for the PFAS in the plots, and each distinct homologous series separated into different KMD-CF_2_ values depending on the composition of their headgroup (left side of **Figure 3A**). To further illustrate this separation, 3D plots of the KMD-CF_2_, *m/z* and CCS values are shown for PFSA, PFCA and FTSA subclasses on the right side of **Figure 3A**. KMD-CH_2_ analyses were performed next to understand if any relationships existed for the studied datasets. As illustrated in **Figure 3B**, the PFAS separated into two lines with a distinct slope in the KMD-CH_2_ versus *m/z* plots. Finally, the mass defect evaluations showed very similar plots to the KMD-CF_2_ analyses since the KMD-CF_2_ scalar was extremely close to 1, **Equation 2**, however a slight slope was observed for each mass defect homologous series (**Figure 3C**).

**Figure 3.**
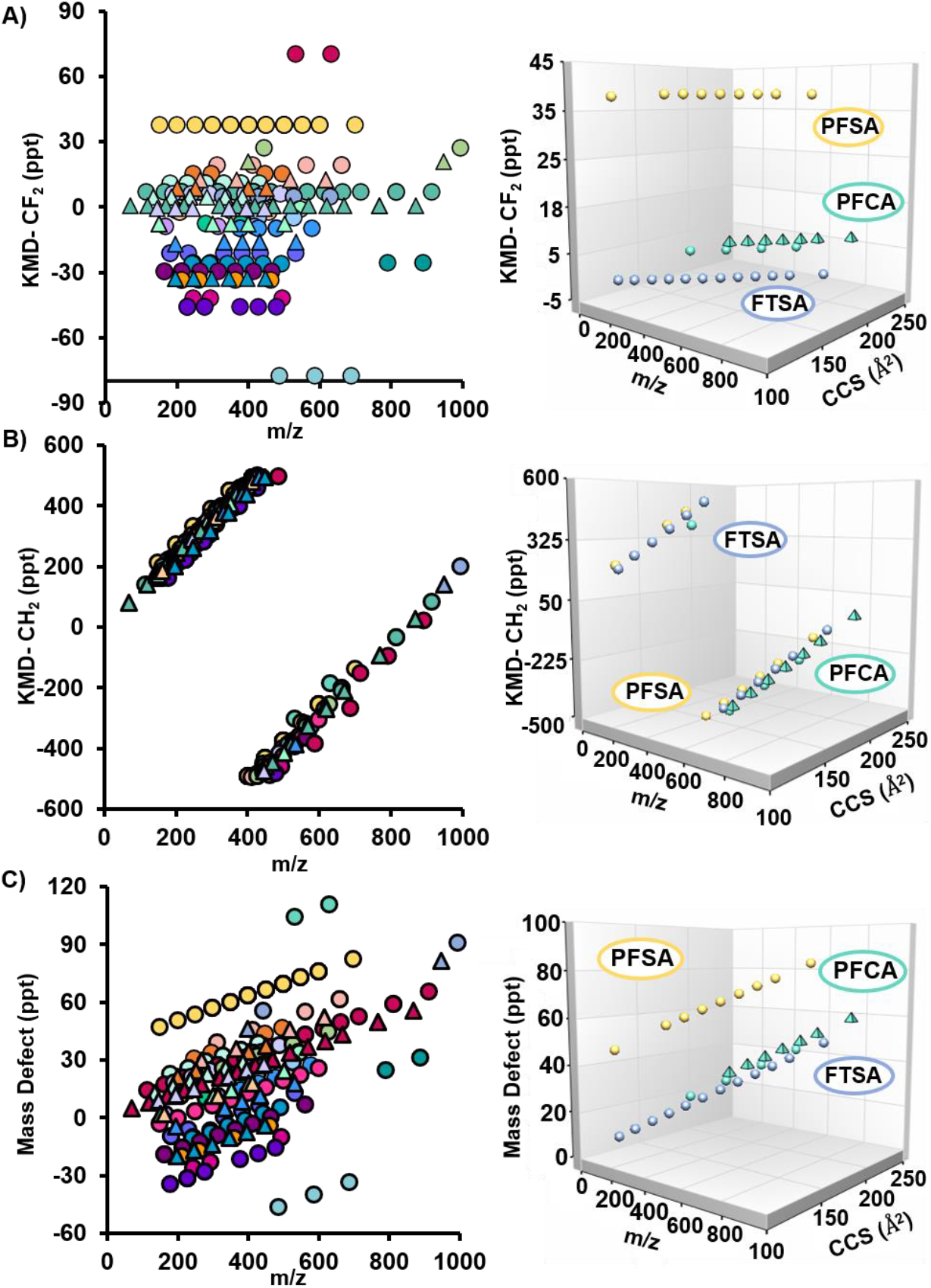
Evaluation of PFAS homologous series. Specific homologous series corresponding to 201 PFAS including PFSA, PFCA and FTSA were assessed by comparing *m/z* values, mass defects, KMDs and CCS values. **A)** In the 2D evaluations of *m/z* versus KMD-CF_2_ (left), the PFAS fall on horizontal lines with a slope of zero. The 3D plots of *m/z*, CCS and KMD-CF_2_ plots (right) also illustrated the flat KMD slope but showcased different CCS values for each PFAS. These relationships were also explored for **B)** KMD-CH_2_ and **C)** mass defect. While two sloped lines were observed for the PFAS with KMD-CH_2_, the mass defect analyses were similar to the KMD-CF_2_ plots but showed a slight slope.

The specific mass defect versus *m/z* relationships observed for the PFAS subclass analyses warranted examination of other molecular types to understand if PFAS were distinct and could be confirmed by the nontargeted analyses. In **Figure 4**, the mass defect and KMDs for PFAS, fatty acids (FAs), bile acids (BAs), pesticides, PCBs, PBDEs, and PAHs were plotted against *m/z*. In the *m/z* versus KMD-CF_2_ plots (**Figure 4A**), the PFAS were all positioned between −100 ppt and 100 ppt on the KMD-CF_2_ axis for the whole mass range examined (**Figure 4A**). Since the other molecules studied did not have a high amount of CF_2_ functional groups, they illustrated different mathematical relationships with some having specific slopes (e.g., PCBs and PBDEs) and others being scattered throughout the graph (e.g., FAs and pesticides). In the *m/z* versus KMD-CH_2_ plots, the two lines for the PFAS intersected with the other molecule types making the small *m/z* value PFAS difficult to distinguish (**Figure 4B**), while the higher *m/z* PFAS were well separated. Finally, in the *m/z* versus mass defect plots, all molecules behaved similar to the KMD-CF_2_ plots as previously observed in the PFAS only studies (**Figure 4C**). Overall, the KMD and mass defect molecular evaluations showed KMD-CF_2_ and KMD-CH_2_ had the most orthogonality and potential for adding confidence to PFAS annotations, while mass defect and KMD-CF_2_ values greatly overlapped and did not provide additional confidence for the molecular identifications. Furthermore, the unique trend lines observed in these plots and the CCS versus *m/z* analyses illustrated their potential for distinguishing PFAS. However, since only a few hundred PFAS chemical standards are available and the EPA estimates over 9,000 PFAS exist due to metabolism and degradation, this lack of standards greatly limits identification capabilities if only experimental values are utilized. In this context, the prediction of CCS values is essential for aiding in the identification of features that are suspected to be new PFAS from mass defect filtering and accurate mass measurements.

**Figure 4.**
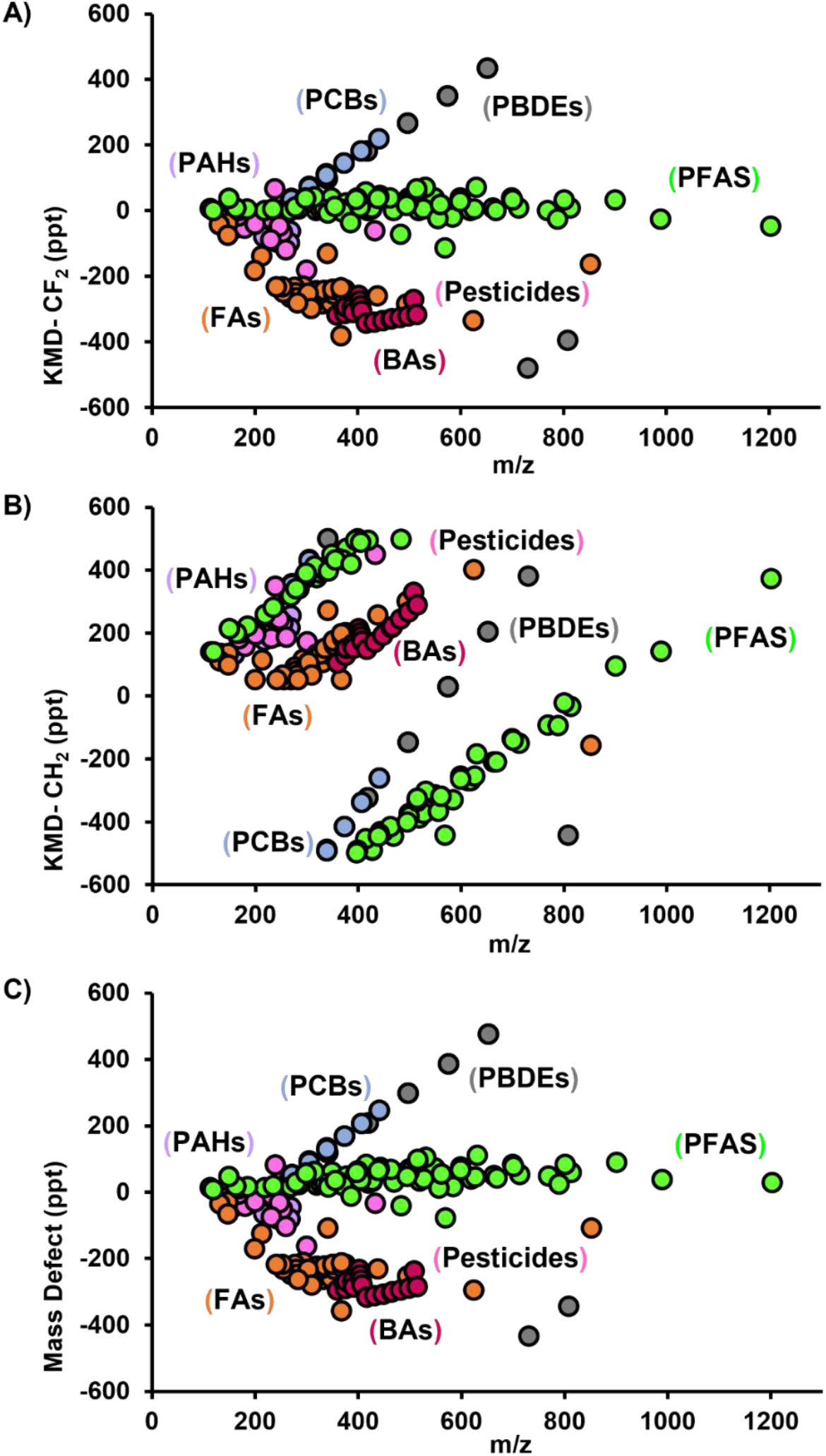
KMD and mass defect evaluations of different molecule types. KMD and mass defect separate PFAS from other molecular classes when plotted against *m/z*. **A)** In the *m/z* versus KMD-CF_2_ plots, all 88 PFAS standards studied were observed between −100 and 100 ppt on the KMD-CF_2_ axis, while other molecular types had different mathematical relationships. **B)** In the *m/z* versus KMD-CH_2_ plots, the PFAS have a distinct slope and **C)** in the mass defect versus *m/z* plots, the PFAS behaved similar to the KMD-CF_2_ plots.

### CCS Prediction for Xenobiotics via Machine Learning

A number of quantum mechanical and ML approaches have been reported to predict CCS values, but xenobiotic applications have been largely lacking in this area.^48,49,53–57^ Though accurate *m/z* and various types of KMD analyses can be used to narrow down chemical classes, many isobaric or isomeric species still exist within the same class. For example, the CompTox database^7^ lists 39 PFAS with the same chemical formula of C_8_HF_15_O_2_, therefore indistinguishable by either KMD or accurate mass analysis. Although tandem MS experiments can potentially distinguish some of these species, many chemically related compounds exhibit similar fragmentation patterns or are present in such low concentrations that even MS/MS analysis is not feasible. While experimentally measured CCS values could be used to filter false positives by matching against a database, CCS databases tend to be largely incomplete due to the lack of available standards. Thus, ML prediction of CCS values is an essential tool in filling in for the lack of standards and providing a way to filter out false positive matches during structural assignments.

Our implementation of ML CCS predictions relies on a linear support vector regression (SVR) model, first trained on a set of known xenobiotic CCS values using molecular descriptors as the input (**Figure S1**). These molecular descriptors are functions that accept molecular identifiers as an input and output numerical data such as molecular weight and volume, atom and ring counts, and average atomic electronegativity. Molecular descriptors are used to predict CCS values *via* multiple ML algorithms including partial least squares regression (PLSR)^48–50^, SVR, and artificial neural networks (ANN).^57^ Once a ML model is accurately trained with low errors, it is used to compute predicted CCS values for all possible xenobiotic candidate structures matching accurate mass and KMD values, and filtering out those falling outside experimental CCS value tolerances.

To evaluate the accuracy of predicted xenobiotic CCS values, PFAS, PBDEs, PCBs and PAHs were studied. Due to similarities in structure, PBDEs and PCBs were combined in a PCB/PDBE group. A total of 1001 ML SVR models were created for each group using a different allocation of molecules for the training (calibration) and test (validation) sets. The models producing the median root mean square error of cross-validation (RMSECV) were then selected for further examination (**Figure 5, Tables S4-S6**). The PFAS model yielded a median prediction error of 1.0% with a root mean square error of validation (RMSEV) of 4.6 Å^2^. Of the 50 PFAS in the validation set, 70% of predictions fell within 3% error, and 90% of predictions were within 5% error. However, five predictions were within 8% error, consisting of four fluoroether species and one cycloalkane. These errors were explained by the steric hindrance of bulky fluorinated alkyl substituents causing systematic overprediction of fluoroethers, and the lack of both fluoroethers and cyclic compounds in the PFAS training set leading to poorer predictions. In the PBDE/PCB SVR model, a median prediction error of 0.49% and a RMSEV of 1.5 Å^2^ were observed. Of the 36 validation compound predictions, 75% fell within 1% error, 92% were within 2% error, and all predictions were within 3% error. Finally, the median prediction error and RMSEV for the PAH validation set were 0.79% and 2.5 Å^2^, respectively. Of the 15 PAH validation predictions, 93% fell within 3% error and all predictions were within 4% error. The only xenobiotic standard outside of the 3% error range was coronene, likely due to its large planar structure that induces enhanced aromaticity. Overall, the prediction errors of ≤1% for all three SVR models compared favorably to other ML-based CCS prediction methods.^48,49,55,57^ For example, the MetCCS SVR approach using a radial basis function has been reported to have median prediction errors of ∼3%;^55^ a deep learning model produced predictions with median errors of 2.6%,^57^ and a partial least squares approach predicted CCS values with median errors close to 2%.^48,49^

**Figure 5.**
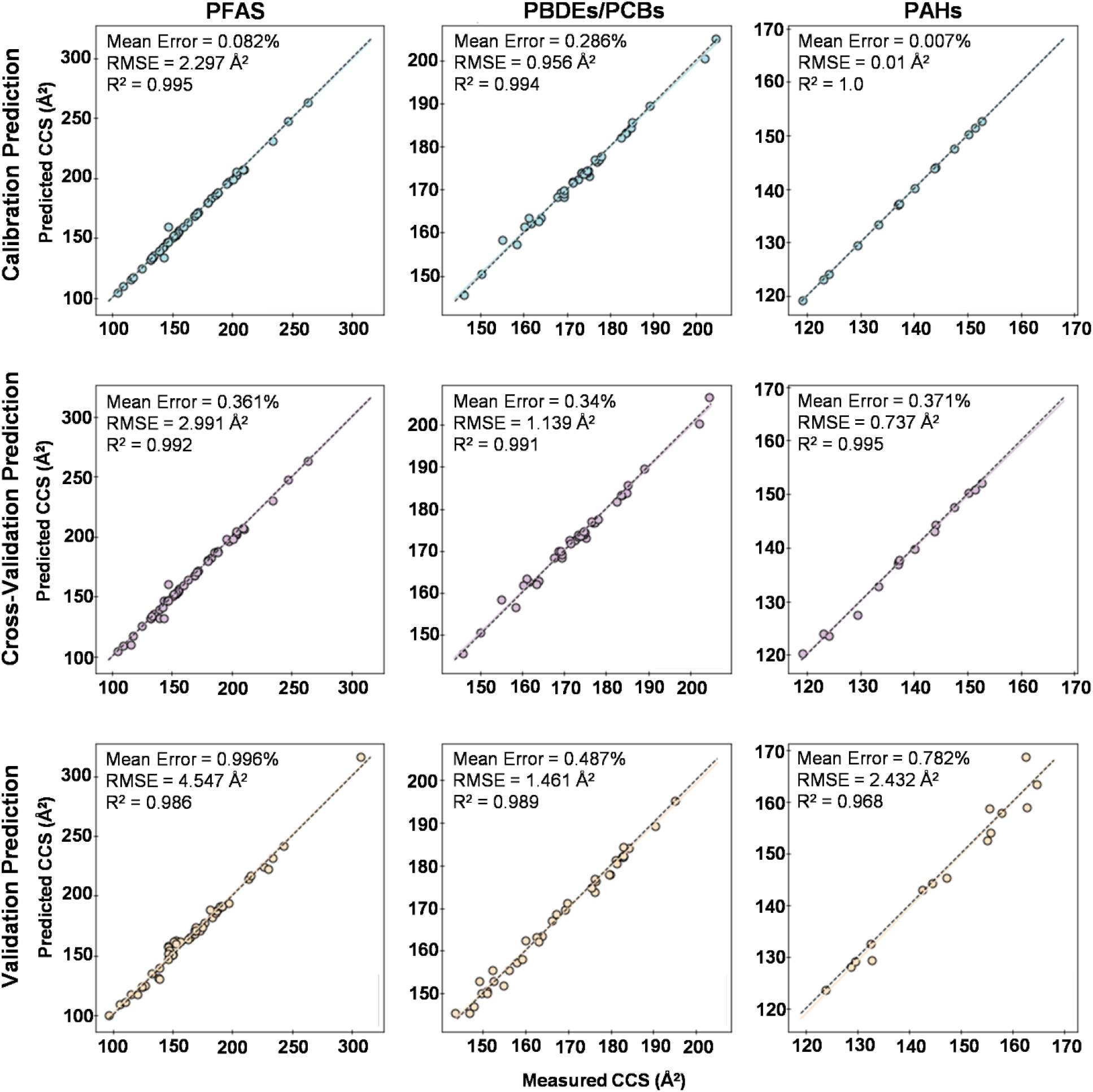
CCS predictions for PFAS, PBDEs/PCBs and PAHs using CCSP 2.0. Three sets of xenobiotics including PFAS (*n* = 102), a combination of PBDEs/PCBs (*n* = 72), and PAHs (*n* = 30) were evaluated with CCSP 2.0. Predictions were made for each set 1001 times with each iteration randomly allocating 50% of the molecules to calibration and 50% to validation. The allocation producing the median root mean squared error (RMSE) of cross-validation is illustrated for each set in their calibration, cross-validation, and validation studies. Calibration and validation points are shown by markers and the ML and 1:1 fits are shown by the colored and dotted lines.

The ML modeling applied here also gives insight into which molecular descriptors are most critical for predictions of each xenobiotic class. Because RFE selects different subsets of molecular descriptors when the train/test allocation is varied, the conservation of descriptors across different allocations indicates their importance in the CCS prediction of a xenobiotic class. The most conserved molecular descriptor classes across these datasets included distance matrices, Barysz matrices, and autocorrelations in topological structure (**Table S7**). Each descriptor class also utilizes information about the relative positions of atoms or sites of unsaturation to describe molecules, enabling this ML approach to produce accurate CCS predictions even among constitutional isomers. Furthermore, the conserved molecular descriptors often utilized weighing schemes based on electronegativity, ionizability and volume, rather than simply mass or elemental formula, which is identical for isomeric species.

The high accuracy of our ML approach combined with its potential to discriminate isobaric species makes it a powerful tool for the structural identification of xenobiotics in the absence of standards. Furthermore, CCS predictions can help detect false discoveries based on exact mass alone. To aid in the annotation of PFAS species not measured in our Skyline library, we used CCSP 2.0 to populate a database of predicted CCS values for a subset of the CompTox PFAS Master List (https://comptox.epa.gov/dashboard/chemical_lists/pfasmaster). The predicted database includes 6138 [Cofta-Woerpel, #750]^-^ and 506 [M-COOH]^-^ CCS predictions (**Tables S8 and S9**). The molecules used to build these models are listed in **Table S4** and **Table S10**, while the model evaluations are available in **Figure S2** and **Figure S3**.

### Xenobiotic Selection Workflow

The potential to greatly reduce nontargeted metabolomic feature lists using ML CCS predictions and mass defect filtering prompted us to develop a comprehensive xenobiotic structural annotation workflow. In this workflow, feature information from nontargeted experimental analyses including CCS, *m/z*, KMD-CF_2_, KMD-CH_2_ and mass defect values, were utilized to narrow down thousands of features to those specifically matching xenobiotic properties. Additional information such as GC or LC retention time and fragmentation can also be incorporated at the end of the workflow to increase identification confidence (**Figure 6**). To showcase the application of this xenobiotic annotation workflow, we chose PFAS due to their important toxicological properties and fluorinated structural properties. The workflow was then implemented as follows. First, following instrumental analysis and creation of the feature list through peak alignment and detection, mass defect filtering was utilized to focus on features belonging to the specific class or classes of interest (**Figure 6A.1)**. Here, specific tolerances for the mass defect, KMD-CF_2_ and/or KMD-CH_2_ are set to either include or remove a feature from the list of candidates. Since PFAS mainly occur between −100 ppt and 100 ppt KMD-CF_2_ (**Figure 6B**) and have specific slopes in the KMD-CH_2_ versus *m/z* plots, tolerances can be set to remove other molecular classes. Additional class filtering can then be achieved by utilizing subclass-specific relationships between CCS and *m/z*, based on either experimental or ML-predicted CCS values (**Figure 6A.2** and **Figure 6C**). Finally, the set of target features is evaluated with KMD-CF_2_ homologous series to determine their formula (**Figure 6A.3**) with further validation using GC or LC dimensions and fragmentation if necessary.

**Figure 6.**
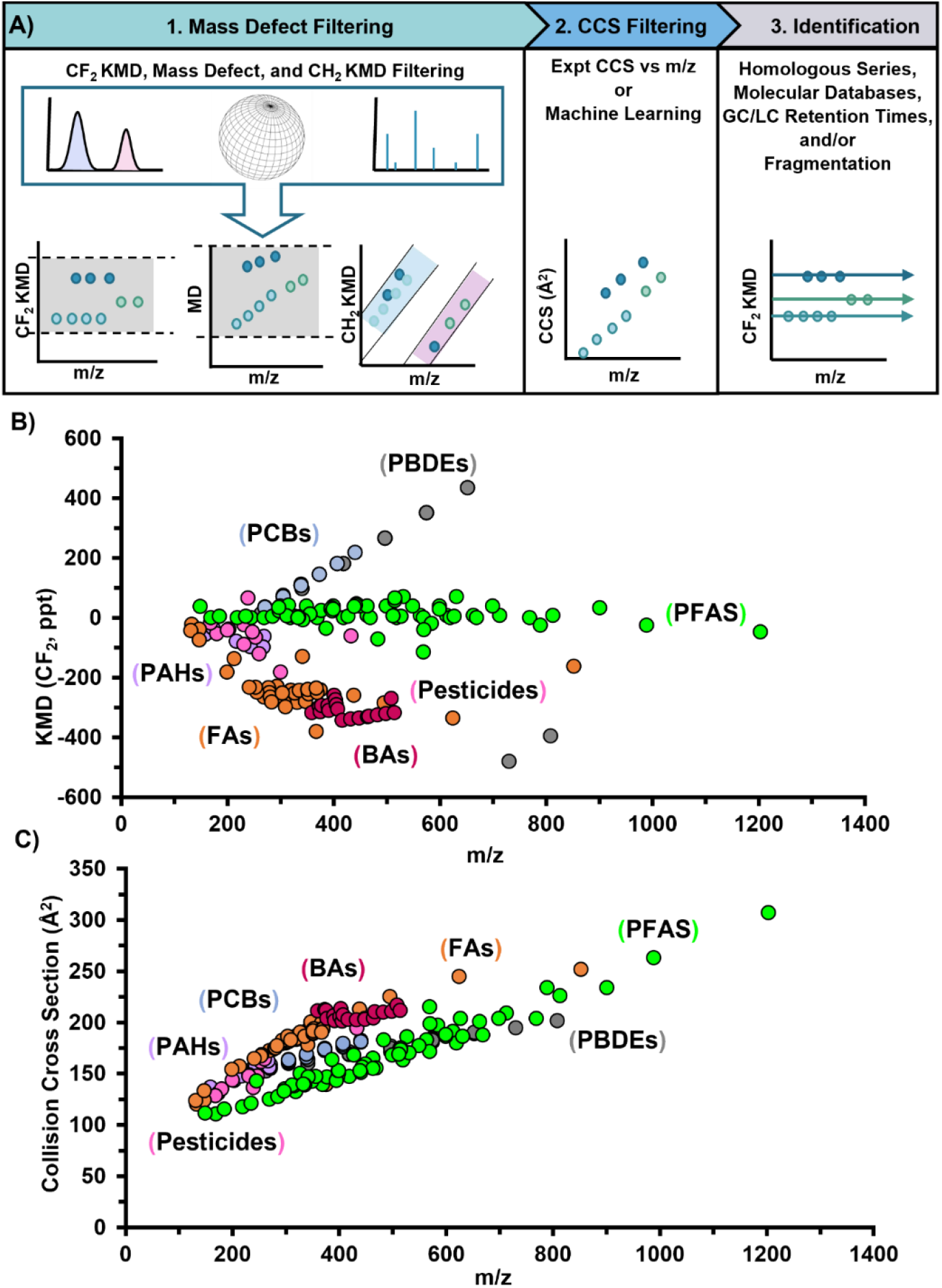
Xenobiotic structural annotation workflow applied to serum. The xenobiotic structural annotation workflow includes A.1) mass defect filtering to discriminate molecular species and homologous series, A.2) CCS filtering with either experimental or ML values to determine class and subclass matches, and A.3) homologous series molecular annotations. To illustrate how selective the first two steps are for PFAS, KMD-CF_2_ filtering based on A.1 is shown in B) and CCS filtering (A.2) is shown in C). All collision cross sections are ^DT^CCS_N2_.

To evaluate the xenobiotic selection workflow, it was applied to a PFAS-specific metabolomic extract from NIST human serum (909c). Since PFAS have been observed in the blood of over 98% of Americans, it was expected that PFAS would be found in this specific reference serum sample.^58,59^ After evaluation with the LC-IMS-MS platform, the resulting serum dataset was evaluated with our Skyline targeted PFAS database containing LC, IMS and MS information. The raw dataset is available on MassIVE (https://massive.ucsd.edu/) with the accession number of MSV000088215. In the targeted data analysis, 17 PFAS were detected at levels significantly higher than those of the blank. These PFAS included 6 perfluorocarboxylic acids, 7 perfluorosulfonic acids, 2 fluorotelomers, 1 perfluoroethercarboxylic acid and 1 perfluoroethersulfonic acid (**Table S11**). Upon comparison with other studies in serum, we observed 7 PFAS that have also been noted in multiple serum studies. These PFAS include perfluoroheptanoic acid (PFHpA), perfluorooctanoic acid (PFOA), perfluorononanoic acid (PFNA), perfluorodecanoic acid (PFDA), perfluoroundecanoic acid (PFUnDA), perfluorohexane sulfonate (PFHxS), and perfluorooctane sulfonate (PFOS).^60^ We were also able to identify 10 additional species in the targeted analyses, which may result from the serum or could be coming from the NIST serum storage conditions as PFAS contaminants are common in vials and storage containers.

The complete set of nontargeted LC-IMS-MS features were next assessed with our annotation workflow to determine additional PFAS not included in our targeted Skyline library. A total of 2,423 LC-IMS-MS features having an absolute abundance greater than 500 were illustrated with Mass Profiler 10.0. This number was then reduced to 129 potential PFAS features using the various mass defect filtering steps described in **Figure 6A.1** as features with mass defects exceeding ±100 ppt or having KMD-CF_2_ exceeding ±80 ppt were discarded. Of the remaining features, those located within two specific regions of a KMD-CH_2_ versus *m/z* plot were investigated; the first region (between 0 Da and 550 Da) included features in the area bounded by *Y* = 1.2018*X* + 39.24 and *Y* = 1.2018*X* − 63.18, while the second region (between 450 Da and 1000 Da) was bounded by *Y* = 1.173*X* − 948.43 and *Y* = 1.173*X* − 1027.92. The 129 potential PFAS features were then further reduced to 80 using the CCS versus *m/z* plot described in **Figure 6A.2**, where features above *Y* = 0.1751*X* + 92.958 were discarded.

To evaluate the efficacy of the filtering steps to select potential PFAS, the 80 remaining features were compared to the subset of the CompTox PFAS Master List selected for CCS prediction. In this comparison, 48 features had masses that matched at least one Master List entry within an error of 20 ppm (**Table S12**). To increase our confidence in the number of PFAS features discovered, the experimental CCS of each feature was compared to the experimentally measured or ML-predicted CCS values of their annotation candidates. Of the 48 features with candidate matches based on mass, only 30 have CCS values within 4% of their candidates. These observations suggest that, though the features may be PFAS, their exact structural isomer may not be present in the database. Thus, additional work utilizing NMR or novel fragmentation approaches such as ultraviolet photodissociation (UVPD) and electron activated dissociation (EAD) is needed to validate potential isomers. Challenges however still occur in these fragmentation analyses as they favor positive mode ionization and PFAS are mainly observed in negative mode.

To help assign structures for the remaining features, we utilized automated CF_2_-homologous series analyses. Here a series was defined as any set of two or more molecules with KMD values matching within 0.0005 and with masses that differ by multiples of CF_2_ (also within 0.0005 Da error). Sixteen serum features matched homologous series identified within the EPA Priority List and PFAS Master List, though most matched entries in the lists and did not require the homologous series analysis. Three previously unannotated features belonged to homologous series (**Table S13**), allowing tentative chemical formula annotation of C_6_F_6_O, C_11_H_5_ClF_14_O_2_ and C_14_H_18_F_11_NO_4_S_2_. The potential chemical formula of C_14_H_18_F_11_NO_4_S_2_ was of particular interest as it had a mass of 536.0431 Da and KMD-CF_2_ of −77.34 ppt. These values placed it within the range of the fluorotelomer thioether amido sulfonic acid (FtTAoS) homologous series (KMD-CF_2_ of ≈ −77.1 ppt). Though this mass does not match any PFAS in the EPA Priority List, it falls directly between 4:2 FtTAoS and 6:2 FtTAoS. The feature’s mass also matches that of 5:2 FtTAoS within 1.5 ppm and its experimental CCS deviates from the predicted CCS of 5:2 FtTAoS by 1.7%, supporting this annotation. The potential presence of this molecule in serum is however particularly interesting as FtTAoS are typically observed with an even number of CF_2_ repeats due to their synthesis protocols, however metabolism could cleave a single CF_2_ unit. While this approach is providing new information of possible PFAS, one caveat to using the homologous series, is that it does not provide information for any PFAS not containing CF_2_ repeat units. Therefore, further investigations are taking place for the other features that survived all filtering steps except the homologous series to determine potential new species of possible interest to the EPA.

## Conclusion

In this manuscript, we evaluated the ability of IMS CCS values and mass defect filtering techniques in a xenobiotic annotation workflow to narrow large metabolomic feature lists to uncover detected xenobiotics of interest. Due to recent connections between exposure and significant health impacts, PFAS were initially studied with the workflow, but it was also extended to PAHs, PCBs, PDBEs and pesticides to show its utility. Initially, we analyzed 88 PFAS standards using LC-IMS-MS to develop a Skyline library and values for ML since very few CCS values for PFAS are currently available. The experimental PFAS CCS and *m/z* values were then compared to other biomolecule and xenobiotic classes, illustrating clear differentiation between the biomolecules and the halogenated xenobiotics. ML was then applied to predict CCS values for PFAS, PCBs, PBDEs and PAHs due to the lack of numerous standards for these xenobiotics. Median errors of ≤1.0% were illustrated for all the xenobiotics CCS values derived from ML when compared to their experimental values. The xenobiotic annotation workflow containing experimental and theoretical CCS values and mass defects was then applied to features obtained for the nontargeted analysis of NIST serum to discover potential PFAS. From the 2,423 metabolomic LC-IMS-MS detected features, the targeted xenobiotic workflow identified 17 PFAS and the nontargeted workflow illustrated 48 potential unknown PFAS candidates. Many of these candidates are being investigated, but one feature matched the formula and ML CCS value for 5:2 FtTAoS, which is currently not on the EPA Priority List but could be a PFAS of future interest. Since this workflow is also applicable to xenobiotics such as PCBs, PBDEs, and PAHs, it shows great promise in targeting other xenobiotics as industries and manufacturers continue to release new chemicals, and the need to evaluate their presence and toxicity, often without standards, is imperative.

## Acknowledgements

This work was funded, in part, by grants from the National Institutes of Health (P30 ES025128, P42 ES027704 and P42 ES031009) and a cooperative agreement with the United States Environmental Protection Agency (STAR RD 84003201). FMF additionally acknowledges support by 1R01CA218664-01 and NSF MRI CHE-1726528. The views expressed in this manuscript do not reflect those of the funding agencies. The use of specific commercial products in this work do not constitute endorsement by the authors or the funding agencies.

## Author Contributions

M.F. and J.N.D. measured the PFAS CCS values with LC-IMS-MS and performed the NIST serum instrumental and targeted data analysis. M.R. and C.W. developed and optimized the machine learning algorithms for PFAS, PCBs, PAHs, and PBDEs. F.M.F. and E.S.B. supervised the machine learning and experimental evaluations and M.F., M.R., F.M.F. and E.S.B. wrote the manuscript.

## Competing Interests

The authors declare no competing interests.

## Notes

https://panoramaweb.org/NCSU%20-%20Baker%20Lab/PFAS%20Library/project-begin.view

### Competing Interest Statement

The authors have declared no competing interest.

