## Supplemental Figures for "Uncovering Xenobiotics in the Dark Metabolome using Ion Mobility Spectrometry, Mass Defect Analysis and Machine Learning"

**This PDF file includes:**

Figs. S1 to S3

References

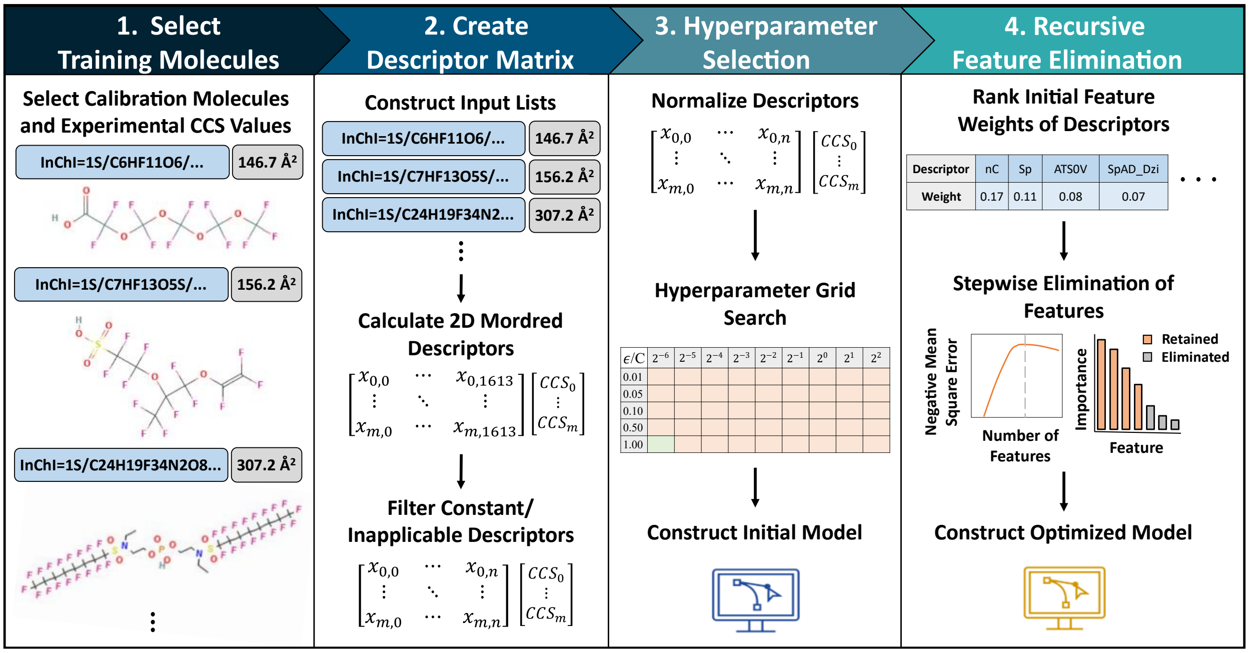


**Figure S1**. Schematic illustration of the CCSP 2.0 workflow for machine learning CCS predictions.

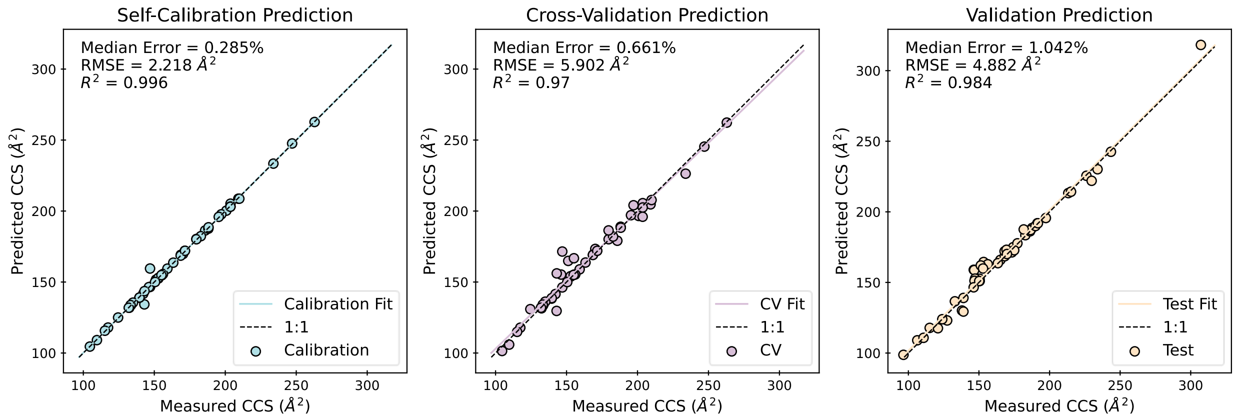


**Figure S2**: CCSP 2.0 was used to predict the collision cross sections of 6138 PFAS in the $[M\text{-}H]^-$ form. Fifty standards were used for calibration and fifty were used for model validation (**Table S3**). Molecular descriptors with outliers (*z*-score > 100) in the application set were discarded. Calibration and validation points are shown by markers and the ML and 1:1 fits are shown by the colored and dotted lines.

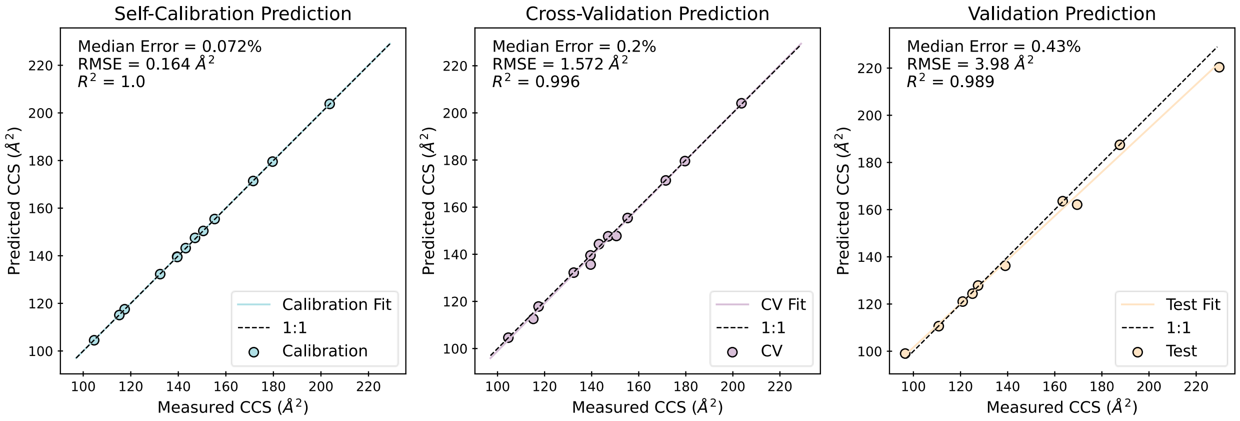


**Figure S3**: CCSP 2.0 was used to predict the collision cross sections of 506 PFAS in the $[M\text{-}COOH]^-$ form. Thirteen standards were used for calibration and ten were used for model validation (**Table Sd**). Molecular descriptors with outliers (*z*-score > 100) in the application set were discarded. Calibration and validation points are shown by markers and the ML and 1:1 fits are shown by the colored and dotted lines.
